# The individual functional connectome is unique and stable over months to years

**DOI:** 10.1101/238113

**Authors:** Corey Horien, Xilin Shen, Dustin Scheinost, R. Todd Constable

## Abstract

Functional connectomes computed from fMRI provide a means to characterize individual differences in the patterns of BOLD synchronization across regions of the entire brain. Using four resting-state fMRI datasets with a wide range of ages, we show that individual differences of the functional connectome are stable across three months to three years. Medial frontal and frontoparietal networks appear to be both unique and stable, resulting in high ID rates, as did a combination of these two networks. We conduct analyses demonstrating that these results are not driven by head motion. We also show that the edges demonstrating the most individualized features tend to connect nodes in the frontal and parietal cortices, while edges contributing the least tend to connect cross-hemispheric homologs. Our results demonstrate that the functional connectome is stable across years and is not an idiosyncratic aspect of a specific dataset, but rather reflects stable individual differences in the functional connectivity of the brain.

**Research highlights:**

- Whole-brain functional connectivity profiles obtained from four resting-state fMRI datasets are unique and stable across 3 months-3 years in adolescents, young adults, and older adults
- Medial frontal and frontoparietal networks tended to be both unique and stable
- Individual edges in the frontal and parietal cortices tended to be most discriminative of individual subjects

## Introduction

Using fMRI and functional connectivity analyses, it is possible to establish a functional connectome for an individual. It has previously been shown that young adults’ functional connectomes are unique and stable across multiple days to one week (Finn et al., 2015; Miranda-Dominguez et al., 2014). This uniqueness and stability can be measured using an identification (ID) test, to ID an individual from a pool of other individuals. To determine the ID rate of a dataset requires at least two test-retest scans from an individual, in which one is designated as a reference scan, and the other a target scan. Using this framework, the ability to identify individuals via their functional connectomes from data acquired in sessions separated by multiple days has been replicated many times in adult subjects (Amico and Goni, 2018; Biazoli et al., 2017; Finn et al., 2017; Finn et al., 2015; Horien et al., 2018; Kaufmann et al., 2018; Noble et al., 2017; Vanderwal et al., 2017; Waller et al., 2017) and in children and adolescents (Kaufmann et al., 2018; Kaufmann et al., 2017). In addition, these studies have consistently shown that regions in the frontal and parietal cortices (i.e. medial frontal and frontoparietal networks) to be important for defining individual uniqueness in functional connectivity data (Amico and Goni, 2018; Finn et al., 2017; Finn et al., 2015; Kaufmann et al., 2017; Miranda-Dominguez et al., 2014; Vanderwal et al., 2017; Waller et al., 2017).

Nevertheless, questions remain as to the stability of the functional connectome over longer periods of time, and the extent to which unique networks remain important in discriminating between individuals over longer time frames, especially given that these networks are comprised of regions that undergo changes throughout the lifespan. For example, it is known that children and adolescents experience marked changes in numerous structural and functional measures, especially in frontal and parietal regions (e.g. Giedd et al., 1999; Gogtay et al., 2004; Gu et al., 2015; Kaufmann et al., 2017; Vasa et al., 2018). Older adults have been shown to exhibit changes in functional connectivity measurements in these regions as well (Chan et al., 2014; Ng et al., 2016; for a recent review, see Damoiseaux, 2017). Hence, these populations provide an intriguing opportunity to test if individual differences in the connectome are stable despite these changes and to determine if unique networks across days remain unique at longer time frames. In fact, a preliminary report using a single dataset suggested that younger subjects (7-15 year olds) do seem to exhibit unique and stable whole-brain connectomes over 1-2 years (Miranda-Dominguez et al., 2018).

Here, using four independent longitudinal fMRI datasets, from four different sites, we set out to confirm these findings and test the hypothesis that the individual functional connectome is stable over a period of years. We extend this work by characterizing the relative importance of specific networks in discriminating between individuals, as well as determining the anatomical locations of the most and least informative edges across years. The findings reveal that connectomes are indeed stable across time spans of at least three years and that the networks driving detection of individual differences across days (i.e. medial frontal and frontoparietal networks) also drive detection across longer time frames.

## Methods

### Overview

We have two main goals in this paper: 1) determine connectome stability using connectome-based identification over the course of years during periods of large-scale changes in the brain and 2) determine the importance of specific networks and anatomical locations in maintaining subject uniqueness (Figure 1). We do this by analyzing four independent resting-state datasets (three comprised of adolescents/younger adults and one comprised of older adults) with multiple scan sessions (i.e. up to three separate time points years apart) and some with multiple test-retest scans per session (i.e. two rest runs per scan session). To aid interpretation we mainly present results from session 1 –session 2 in the main text, but provide results from the other sessions and test-retest scans in the Supplementary Material.

**Figure 1.**
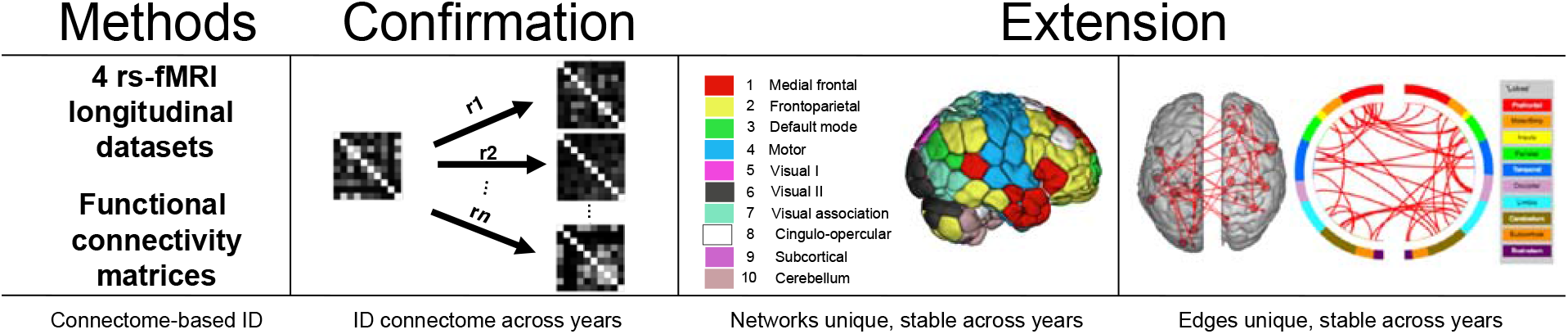
Overview of the goals of this paper. To test connectome-stability, we used connectome-based identification to determine if individual differences in functional connectivity data are detectable across years. The stability and uniqueness of networks across years is also characterized, in addition to determining the locations of the most and least informative edges.

### Description of datasets

We utilized four longitudinal rs-fMRI datasets of neurotypical subjects. The University of Pittsburgh School of Medicine dataset, the University of Utah dataset, and the University of McGill dataset (hereafter referred to as Pitt, Utah, and UM, respectively) were downloaded from the Consortium for Reliability and Reproducibility (CoRR; http://fcon_1000.projects.nitrc.org/indi/CoRR/html/samples.html; Zuo et al., 2014); the Southwest University Longitudinal Imaging Multimodal dataset (hereafter referred to as SLIM; Liu et al., 2017) was downloaded through the International Data-sharing Initiative (INDI; http://fcon_1000.projects.nitrc.org/). All datasets were collected in accordance with the institutional review board or research ethics committee at each site. All resting-state scans were acquired on Siemens 3-T Trio scanners. Relevant demographic characteristics and imaging parameters are described in Table 1; full details of the Pitt, SLIM, and UM datasets can be found elsewhere (Hwang et al., 2013; Liu et al., 2017; Orban et al., 2015). To provide a fuller picture of the demographic data we provide histograms of the number of years between scans and age at the time of the first scan for all datasets (Supplemental Figure 1 and 2). We note that in the Utah data set, two resting-state scans were provided for session 2 (hereafter session2a and session 2b) and with the UM data, two resting-state scans were provided for session 1 (session1a and session1b) and for session 2 (session2a and session2b).

**Table 1.** Demographic and imaging characteristics of datasets used in this study.

|  | **SLIM** | **Pitt** | **UM** | **Utah** |
| --- | --- | --- | --- | --- |
| Subjects in session 1 and 2 (females) | 105 (49) | 93 (45) | 79 (58) | 26 (0) |
| Subjects in session 3 (females) | 105 (49) | 30 (16)<sup>a</sup> | - | - |
| Low motion subjects session 1-session 2 | 47 | 44 | 27 (Session 1a – session 2a) | 16 (Session 1 – session 2a) |
| Low motion subjects session 1-session 3 | 38 | 15 | 27 (Session 1a – session 2b) | 14 (Session 1 – session 2b) |
| Low motion subjects session 2-session 3 | 40 | 16 | 25 (Session 1b – session 2a) | - |
| Low motion subjects session 1b – session 2b | - | - | 25 (Session 1b – session 2b) | - |
| Age at scan 1 | 19.67 ± 0.96 years | 15.23 ± 2.83 years | 65.3 ± 6.28 years | 20.23 ± 8.28 years |

| (mean± SD)<sup>b</sup> | | | | |
| --- | --- | --- | --- | --- |
| Years between scan 1 and 2 (mean± SD) | 0.84 ± 0.29 years | 1.76 ± 0.41 years | 0.305 ± 0.067 years | 2.54 ± 0.29 years |
| Years between scan 2 and 3 (mean± SD) | 1.532± 0.18 years | 1.53 ± 0.30 years | - | - |
| Years between scan 1 and 3 (mean± SD) | 2.37 ± 0.28 years | 3.16 ± 0.26 years | - | - |
| Scan duration in minutes (volumes) | 8 (242) | 5 (200) | 5 min 45 sec (150) | 8 (240) |
| TR in seconds | 2 | 1.5 | 2 | 2 |

a. Scan session 2 and 3 occurred on the same day in one female subject in this session; we removed it when conducting analyses that required us to incorporate the number of years between scans 2 and 3.
b. All datasets provided age in years at scan session 1. For SLIM and Utah, the ages were reported as integers in years only, while the time between scan sessions was reported in days. To standardize the reporting of age among all four datasets, we only report age at session 1 here.

Note that because of numerous differences between the datasets (eyes open/closed during rest, resting-state scan collected after other functional runs, differences in number of years between scans, etc.) we did not compare ID rates between datasets; rather, our focus was on defining the upper bounds of identifiability within each dataset.

### Preprocessing

The preprocessing strategy used has been described in detail elsewhere (Greene et al., 2018). All analyses were performed using BioImage Suite (Joshi et al., 2011) unless otherwise indicated. We note that we only preprocessed the Pitt, Utah, and UM subjects, as the SLIM data was preprocessed beforehand as in Liu et al. (2017); for SLIM, we downloaded (http://fcon_1000.projects.nitrc.org/indi/retro/southwestuni_qiu_index.html) pre-calculated connectivity matrices based on the 268 node functional atlas described below. The preprocessing steps included: skull-stripping the 3D magnetization prepared rapid gradient echo (MPRAGE) images using optiBET (Lutkenhoff et al., 2014) and performing linear and non-linear transformations to warp a 268 node functional atlas from MNI space to single subject space using BioImage Suite as in Greene et al. (2018). Functional images were slice-time and motion corrected using SPM8 (https://www.fil.ion.ucl.ac.uk/spm/software/spm8/). Covariates of no interest were regressed from the data, including linear, quadratic, and cubic drift, a 24 parameter model of motion (Satterthwaite et al., 2013) mean cerebral-spinal fluid signal, mean white matter signal, and the overall global signal. Data were temporally smoothed with a zero-mean unit-variance low-pass Gaussian filter (approximate cutoff frequency of 0.12 Hz). The results of skull-stripping, non-linear, and linear registrations were inspected visually after each step. Subjects were excluded from further analyses in Pitt and SLIM due to incomplete coverage of functional scans (mostly in the cerebellum).

### Node and network definition

We used a 268 node functional atlas described previously (Finn et al., 2015). For each subject the average time-course of each region of interest (node in graph theoretic terminology) was calculated, and the Pearson correlation coefficient was calculated between every other node to achieve a symmetric 268 x 268 matrix of correlation values representing edges (connections between nodes) in graph theoretic terminology. We subsequently normalized the matrix to z-scores via a Fisher transformation and only considered the upper triangle of the matrix, yielding 35,778 unique edges for whole-brain analyses. These nodes were then grouped into 10 functional networks as in previous work (Finn et al., 2015; Noble et al., 2017). Network names are listed in Figure 5a.

In the SLIM and Pitt datasets, even after excluding subjects missing significant amounts of data in the functional scan, many subjects were still missing edges in the whole-brain connectivity matrix due to incomplete coverage. To ensure standardization within a dataset, if any subject was missing an edge, we removed that edge from all subjects. Of the 35,778 edges in the whole-brain functional connectome, 31,626 edges/subject remained in the SLIM dataset; 29,646 edges/subject remained in the Pitt dataset. All 35,778 edges were covered for all subjects in the Utah and UM datasets (after removing 1 subject missing ~1000 edges from all scans in UM). See Supplemental Table 1 for the final list of the subject IDs used in this study. See Supplemental Figure 3 for the proportion of edges remaining in network pairs for SLIM and Pitt.

### Identification procedure

The identification procedure has been described in detail previously (Finn et al., 2015). Briefly, a database is first created consisting of all subjects’ connectivity matrices from a particular session for a specific dataset. In an iterative process, a connectivity matrix from a subject is then selected from a different session and denoted as the target. Pearson correlation coefficients are calculated between the target connectivity matrix and all the matrices in the database. If the highest Pearson correlation coefficient is between the target subject in one session and the same subject in the second session, this would be recorded as a correct identification. We repeat the identification test such that each subject serves as the target subject once. This process is repeated until identifications have been performed for all subjects, sessions, and database-target combinations. We then averaged both database-target pairs (because these can be reversed) for a dataset to achieve an average ID rate for a given scan pair. This approach was used for whole-brain ID as well as network-based ID, in which connectivity matrices are formed using only the edges from a particular network of interest. Statistical significance was assessed via permutation testing, in which subject identities were randomly shuffled and ID was performed with the incorrect labels. To compare ID rates between pairs of networks in a dataset we performed bootstrapping: we randomly subsampled ~80% of the subjects in each iteration and performed identification. The process was repeated 1000 times to generated 95% confidence intervals and determine significance.

### Controlling for motion and artificial elevations in ID rate

To avoid confounds due to motion, we performed identification on low motion subjects (i.e. a mean frame-frame displacement (FFD) threshold of < 0.1mm for both scans) as described previously (Horien et al., 2018); the number of low motion subjects in each dataset is provided in Table 1. Imposing a threshold like this has previously been shown to limit confounds due to motion in different samples (Greene et al., 2018).

To further determine that identification was not due to idiosyncratic head movements specific to an individual across sessions, we conducted identification analyses using estimates of head movement parameters which has been described in detail elsewhere (Finn et al., 2015). Briefly, we calculated discrete motion distribution vectors for each subject based on FFD over an entire scan. We computed the mean and standard deviations of the FFD across all subjects for each dataset. Based on the number of volumes collected, we then specified 15 bins for the Utah and SLIM subjects, 10 bins for the Pitt subjects, and 8 bins for the UM subjects to span the grand mean +/− 3 standard deviations, and motion distribution vectors were subsequently calculated. These vectors were then submitted to the identification procedure. The results verified that motion was not a factor in the identification.

Next we tested to see if the results were driven by an artificially reduced sample size. For this we examined all the subjects and repeated these analyses. In the context of identification this increases the difficulty of a successful ID in two ways: first, by increasing the number of subjects (choices) that a subject may incorrectly match to, and second, by introducing “harder to identify” subjects into the pool (i.e. subjects with a mean FFD > 0.1 mm exhibit less similar functional connectivity patterns from scan to scan and are harder to identify; Amico and Goni, 2018; Horien et al., 2018).

### Edge-based and linear regression analyses

To determine the role of specific edges in the identification process, we performed calculations to quantify highly unique and highly consistent edges via the differential power (DP) measure and the group consistency measure described in detail elsewhere (Finn et al., 2015); we provide a brief overview here. DP provides an estimate, for each given edge, of the likelihood that within-subject similarity is higher than that between subjects. To calculate the DP, the product of edge values from time 1 and time 2 from the same subject is compared to the product of time 1 and time 2 from unmatched subjects. Edges with high DP values are considered helpful in identification. To calculate the group consistency measure, we multiply an edge value from time 1 and time 2 across all edges for all subjects and calculate the mean for each edge. Edges with high values in this measure are therefore high across all individuals in the group and are not helpful in identification. In the course of these analyses, we also determined the network(s) to which any given pair of edges belong (within- and between-networks); to account for differences in network sizes in both of these analyses, we divided the number of significant edges in a network pair by the number of total edges in the network.

To investigate the relationship between within-subject correlation scores (self-correlation), age, and time between scans, we performed linear regression analyses between self-correlation and the number of years between scans; we also correlated self-correlation scores with age. We further performed partial correlation analyses in which we assessed the relationship between self-correlation scores and age while controlling for time between scans and vice-versa.

### Code availability

Analyses were conducted in Matlab; code is available online at https://www.nitrc.org/frs/?group_id=51. Questions regarding using or adapting these scripts should be directed to the corresponding author (CH). The corresponding author assumes all responsibility for accuracy/integrity of all data and code.

## Results

### Whole-brain identification results

Analyses using the whole-brain connectivity matrix and low motion subjects (mean FFD < 0.1 mm for both sessions) yielded identification rates well above chance using all four datasets (P < 0.0001; Figure 2A). To determine if ID rates within a dataset were artificially elevated by the reduced sample sizes introduced by the motion threshold, we performed identification again with all subjects in a given dataset. Note that this increases the difficulty of the ID procedure in two ways: by increasing the number of subjects a participant might incorrectly match to, and by introducing harder to identify subjects into the ID pipeline, as high motion subjects tend to look less similar from scan-to-scan (Amico and Goni 2018), which might decrease ID rates independent of sample size (Horien et al., 2018). Nevertheless, when we included all subjects in the ID pool, we observed ID rates again highly above chance (*P* < 0.0001; Figure 2B).

**Figure 2.**
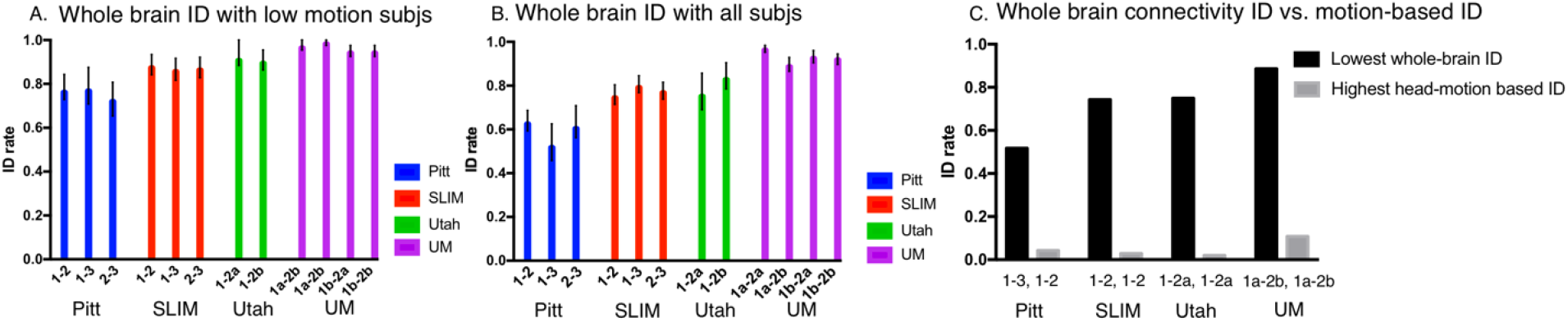
Identification results using whole brain data. Each dataset is indicated below the x-axis, along with the sessions that were involved in identification. ID rate is indicated on the y-axis. Error bars correspond to 95% confidence intervals obtained via bootstrapping. Panel A: results using only low motion subjects – only the subjects with a mean FFD < 0.1 mm were included in this analysis. Panel B: results using all subjects from a dataset – both high and low motion subjects were included. Panel C: For visualization purposes here we are showing the lowest whole-brain ID rate obtained using all subjects (i.e. the lowest rate achieved in Panel B in a given dataset) compared to the highest head motion-based ID from a given dataset. The session pair resulting in a given ID rate is shown below the appropriate bar. (For example, Pitt session 1-session 3 gave the lowest whole-brain ID rate in Panel 1B, while the highest head-motion based ID for Pitt was obtained in session 1-session 2). Note that the lowest whole-brain ID and the highest head motion-based ID do not necessarily have to come from the same scan pair.

Using this full sample, we next determined if subject-specific movement patterns in the scanner were driving the high ID rates. Although based on previous work this is unlikely (Amico and Goni, 2018; Finn et al., 2015; Finn et al., 2017; Horien et al., 2018; Vanderwal et al., 2017), we calculated a motion distribution vector (see Methods for details) capturing patterns of movement for each subject in a scan and used this for identification. The highest ID rates we achieved in each dataset were 1.92%, 2.9%, 4.3%, and 10% for Utah, SLIM, Pitt, and UM, respectively, (Figure 2C), well below the lowest ID rate we achieved using the whole-connectivity matrix, consistent with other identification studies showing motion does not appear to be driving ID. Grouping subjects by gender and performing ID revealed similarly high rates (and no effect of gender on ID), as did performing ID of high motion subjects only (P < 0.0001; Supplemental Figure 4). Thus, we conclude that the functional connectivity matrix is unique and stable at time-frames of three months up to three years, even in individuals undergoing periods of rapid brain development. Importantly, these results do not seem to be driven by motion.

### Within-subject correlation scores, age, and time between scan results

To further investigate connectome stability, we next examined the relationship between self-correlation scores (the correlation of a subject from time 1 to time 2), age, and years in between scans by performing linear regression analyses. To control for motion effects, we imposed the same 0.1 mm mean FFD as above. We observed that there appears to be a negative correlation between self-correlation and time between scans, though the relationship was not statistically significant (all *P* > 0.05 with the exception of SLIM session1-session2, *r* = -0.3105, *P* = 0.0337; and UM session1b-session2a, *r* = -0.5528, *P* = 0.0042; Figure 3). There was no consistent relationship between self-correlation and age, with the only significant result coming from Utah session1-session2a (*r* = -0.6205, *P* = 0.0103; all other *P*-values > 0.05; Figure 4).

**Figure 3.**
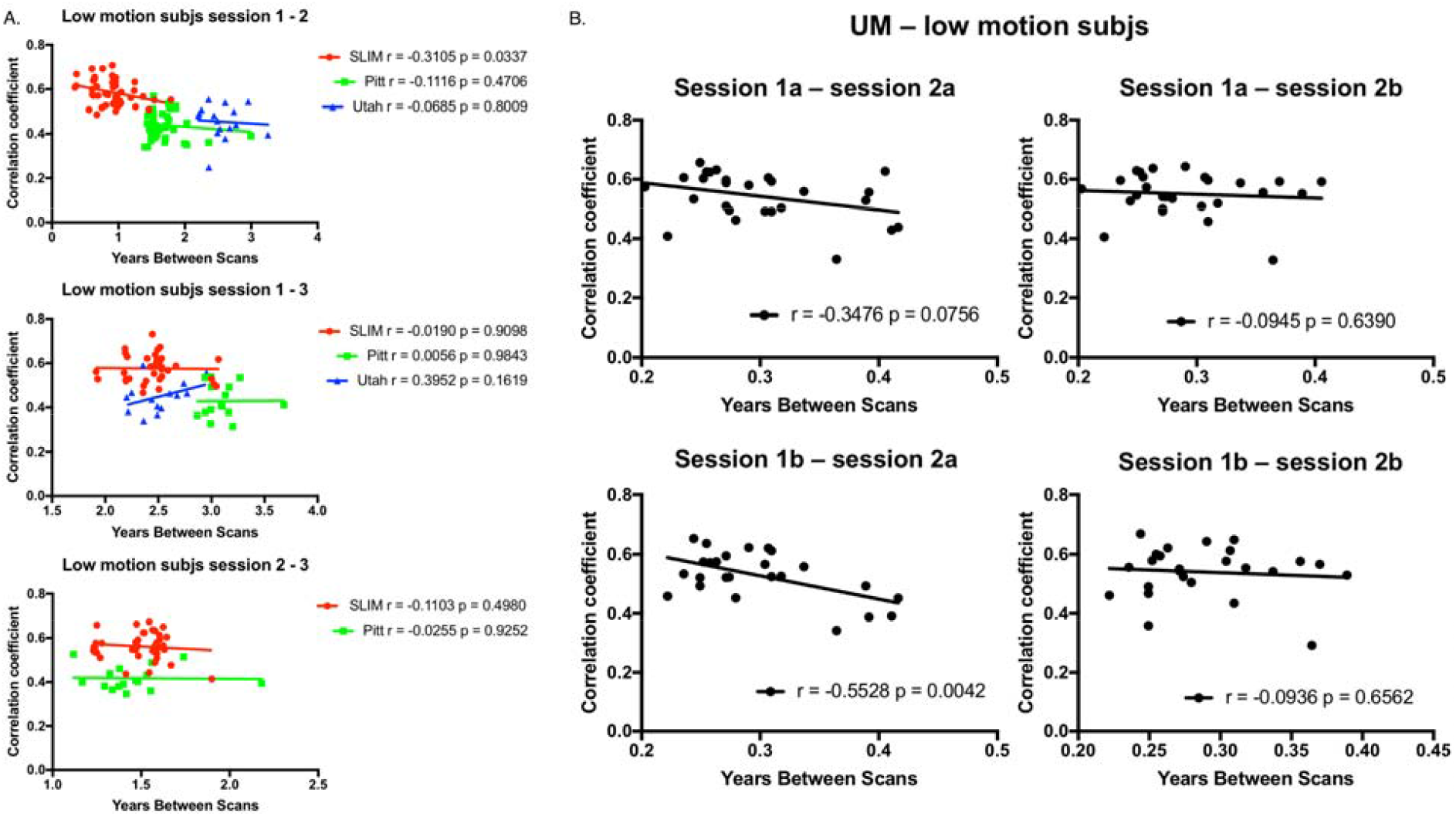
Plotting self-correlation against years in between scans for low motion subjects. Panel A: Results for SLIM, Pitt, and Utah. Panel B: Results for UM. Each dataset is indicated by the appropriate color and symbol. Results of linear regression analyses are shown on the appropriate graph.

**Figure 4.**
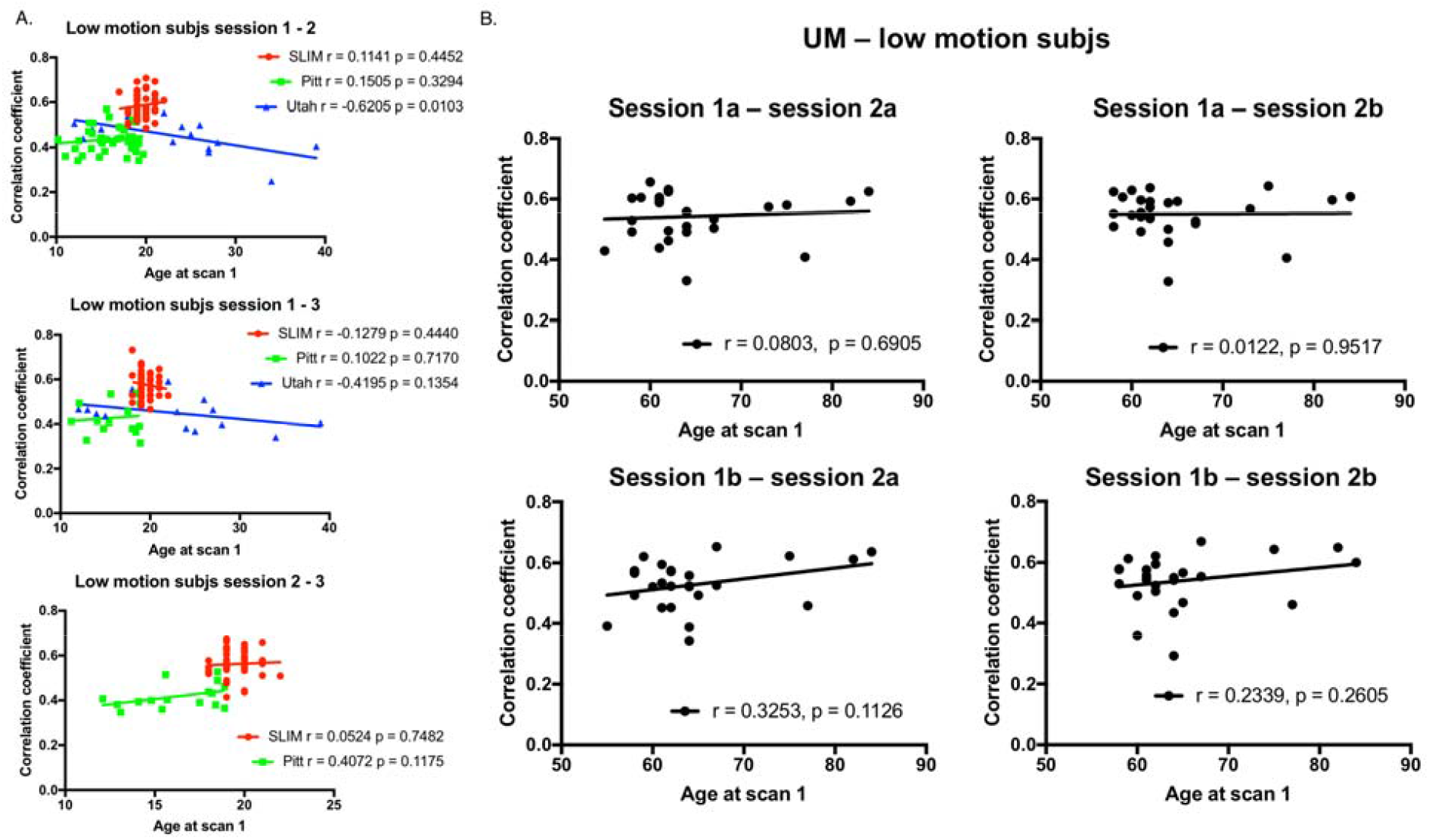
Plotting self-correlation against age at scan one for low motion subjects. Panel A: Results for SLIM, Pitt, and Utah. Panel B: Results for UM. Each dataset is indicated by the appropriate color and symbol. Results of linear regression analyses are shown on the appropriate graph.

We also performed analyses with all subjects in a given dataset, as well as subjects grouped by gender, and with high-motion subjects only (Supplemental Figures 5-12). We observed largely consistent results, in that there is an (expected) negative correlation between self-correlation and time between scans, though the relationship is weak; there is no clear relationship between age and self-correlation score; and motion does not appear to be driving the results (Supplemental Figures 5-8). Further, no clear trends emerged due to gender (Supplemental Figures 9-12).

In addition, we performed partial correlation analyses of the low motion subjects to explore the relationship between self-correlation and years between scans while controlling for age (and vice-versa). No clear patterns emerged (Supplemental Table 2).

Taken together, these data reinforce that self-correlation scores tend to be stable between scans and there is no clear relationship between self-correlation and age. In addition, the results do not seem to be driven by motion.

### Network-based identification results

Having determined whole-brain connectivity data is stable across years, we next tested the contributions of specific networks to this stability. We grouped the connectivity data for each subject into 10 functional networks (Figure 5A), and subsequently performed identification analyses using only the edges from a given network. Given the performance of the medial frontal and frontoparietal networks in previous work (Finn et al., 2015; Waller et al., 2017; Kaufmann et al., 2017), we also combined the edges from these networks (hereafter ‘combined network 1+2’) and performed identification.

**Figure 5.**
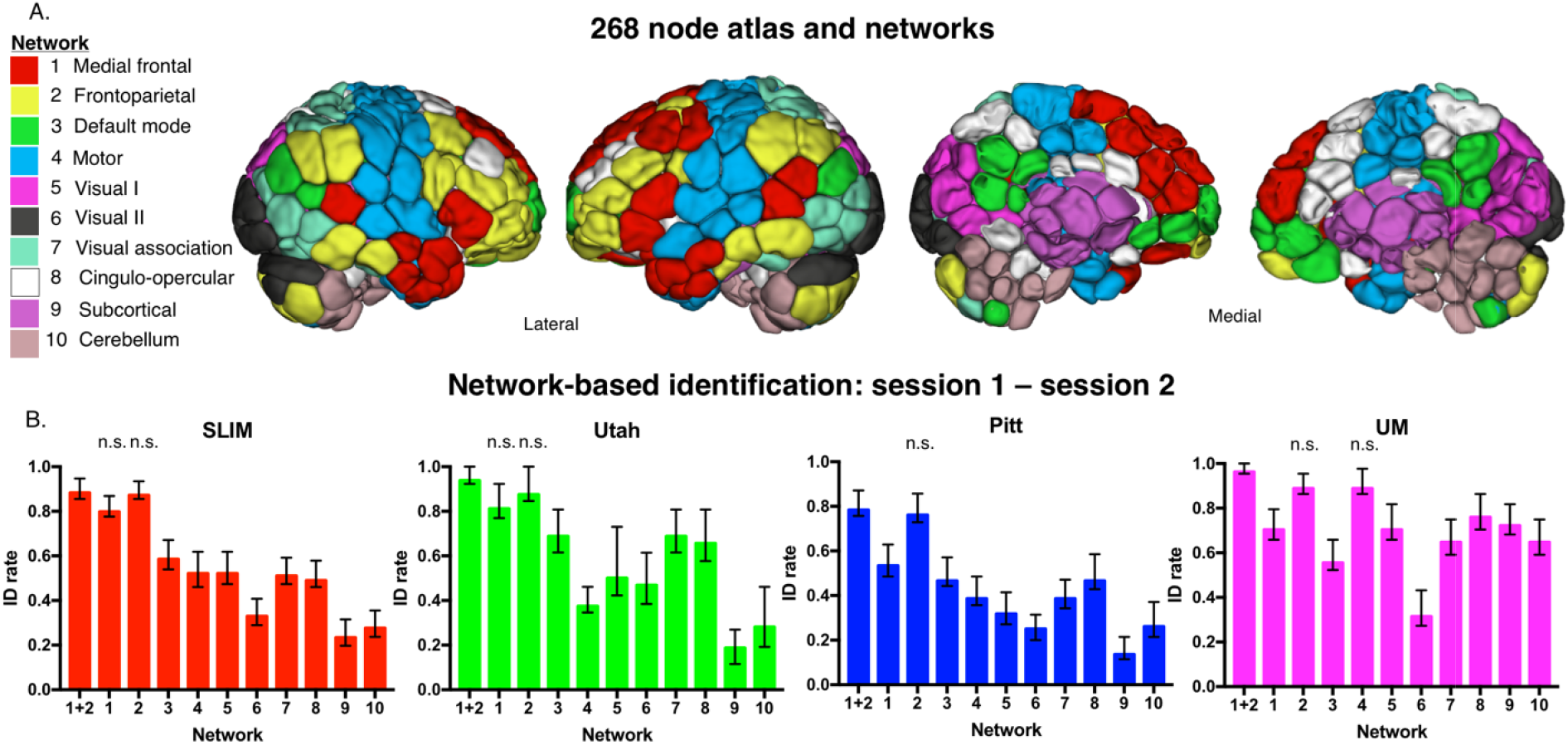
Network-based identification of low motion subjects. Panel A: Node and network labels. We utilized a 268-node functional atlas. Nodes were further grouped into the 10 functional networks indicated here. Network names are to the left; anatomic locations are shown on the brains to the right. Panel B: We performed identification using only within-network edges (networks 1 – 10; indicated below the x-axis of each graph). We also combined networks 1 and 2 and performed identification with these edges (indicated as ‘1+2’ below the x-axis). Dataset name is indicated at the top of the graph. Networks that do not have a statistically lower ID rate (*P* > 0.05) than combined network 1 and 2 are labelled ‘n.s.’ above the corresponding network; all other networks that are unlabeled have an ID rate that is lower than combined network 1 and 2 (*P* < 0.05; i.e. these networks do have ‘n.s’ above the bar). Error bars correspond to 95 percent confidence intervals obtained via bootstrapping. Note that we considered Utah session 2a as session 2; for UM we considered session1a as session 1 and session 2a as session 2.

Focusing on the low motion subjects from session 1 – session 2, we observed that edges in networks 1 and 2 tended to lead to the highest ID rates, as well as combined network 1+2 (Figure 5B), consistent with the original fingerprinting work of Finn et al. (2015). For example, at an alpha of 0.05, combined network 1+2 had higher ID rates than all other networks in SLIM (except compared to individual networks 1 and 2); all other networks in Utah (except compared to individual networks 1 and 2); all other networks in Pitt (except compared to individual network 2); and all other networks in UM (except compared to individual networks 2 and 4). We observed similar patterns of performance when we repeated identification on all subjects in a dataset: networks 1 and 2 resulted in the highest ID rates (Supplemental Figure 13).

To further assess the importance of combined network 1+2 in defining individual uniqueness, we next assessed ID performance relative to between-network pairs of edges in the low motion subjects of session 1 – session 2 (Figure 6). At an alpha of 0.05, we observed that combined network 1+2 resulted in higher ID rates than 43/45 between-network edges in SLIM (non-significant network pairs: 1-2, 5-6); 42/45 between-network edges in Utah (non-significant network pairs: 1-2, 2-7, 2-8); 42/45 between-network edges in Pitt (non-significant network pairs: 1-2, 4-5, 5-6); and 42/45 between-network edges in UM (non-significant network pairs: 1-2, 4-5, 5-6). These results echo the original fingerprinting work of Finn et al. (2015), emphasizing the importance of the medial frontal and frontoparietal network. Performing between-network ID on all subjects in a dataset resulted in a similar pattern: the highest ID rates tended to result from edges connecting to networks 1 and 2 (Supplemental Figure 13).

**Figure 6.**
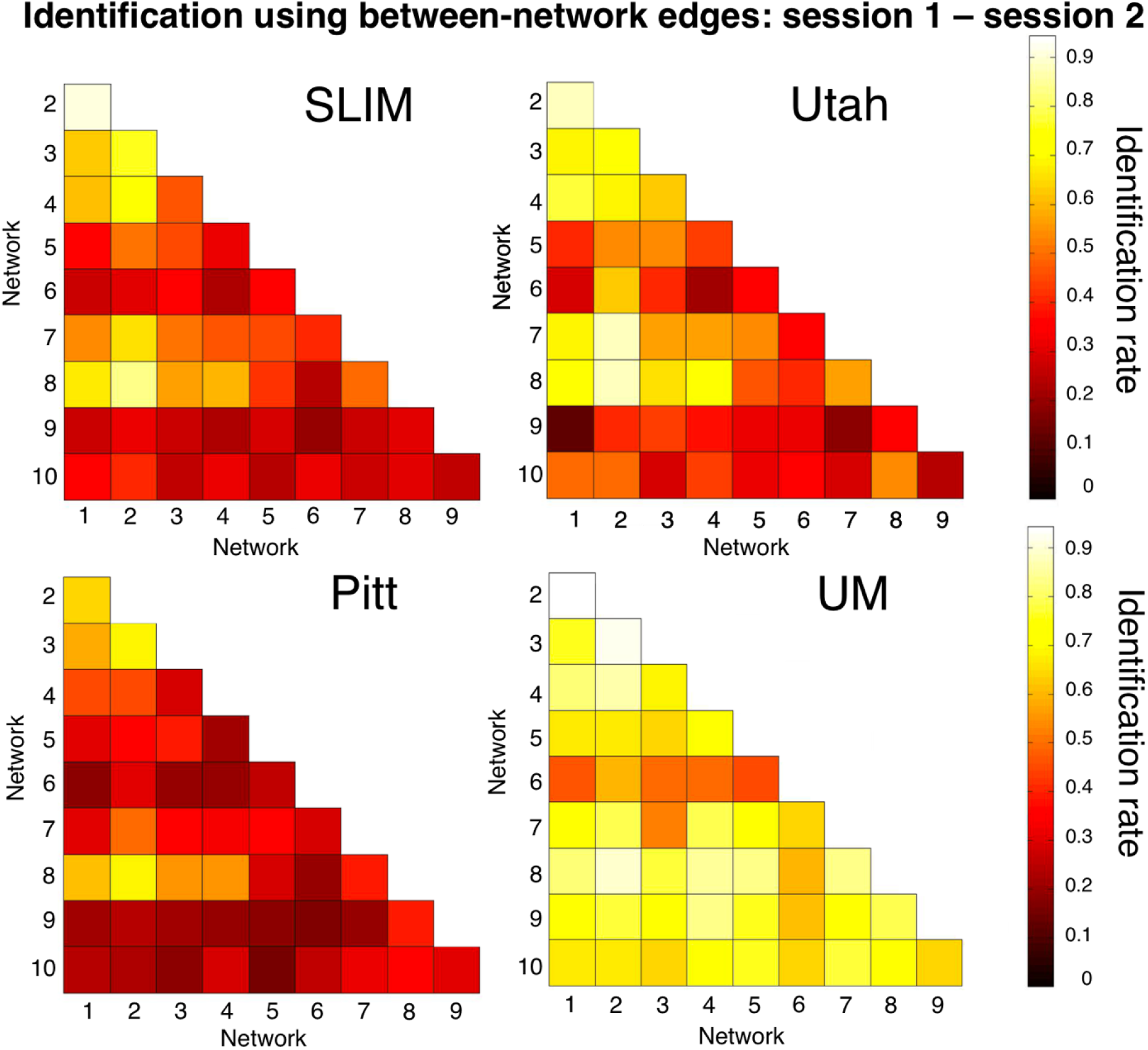
Between-network identification of low motion subjects. We repeated the identification analyses using only the between-network edges. Shown are the results from session 1-session 2. Dataset name is indicated above the appropriate matrix. The scale of the color bars is the same for all matrices. Note that we considered Utah session 2a as session 2; for UM we considered session1a as session 1 and session 2a as session 2. Note also that the diagonal of the matrix (which is not shown) consists of the within-network edges; these data are shown in Figure 5B.

### Edge-based analyses results

Having established that the medial frontal and frontoparietal networks are stable across years in individual subjects, we lastly assessed the importance of specific edges to subject uniqueness in the low motion subjects of session 1 – session 2. Using the DP measure, which calculates how characteristic an edge tends to be, we were able to determine which edges were important in the identification process. For visualization purposes we show the anatomical locations of DP edges in the 99.9 percentile (Figure 7). Significant edges tended to be distributed across the entire brain, with particular representation in prefrontal, temporal, and parietal cortices. Less stringent thresholds resulted in similar overall patterns (albeit with more edges; Supplemental Figure 14). Quantifying network representation revealed that networks 1 and 2 tended to harbor significant edges (Figure 8): approximately 78% of edges were within or connected to networks 1 and 2 in Utah; approximately 60% in SLIM; approximately 58% in Pitt; and approximately 48% in UM. The overrepresentation of significant edges in networks 1 and 2 held at a range of thresholds (Supplemental Table 3). DP results were stable when we reperformed the analyses on all subjects in dataset (Supplemental Figure 15 and 16).

**Figure 7.**
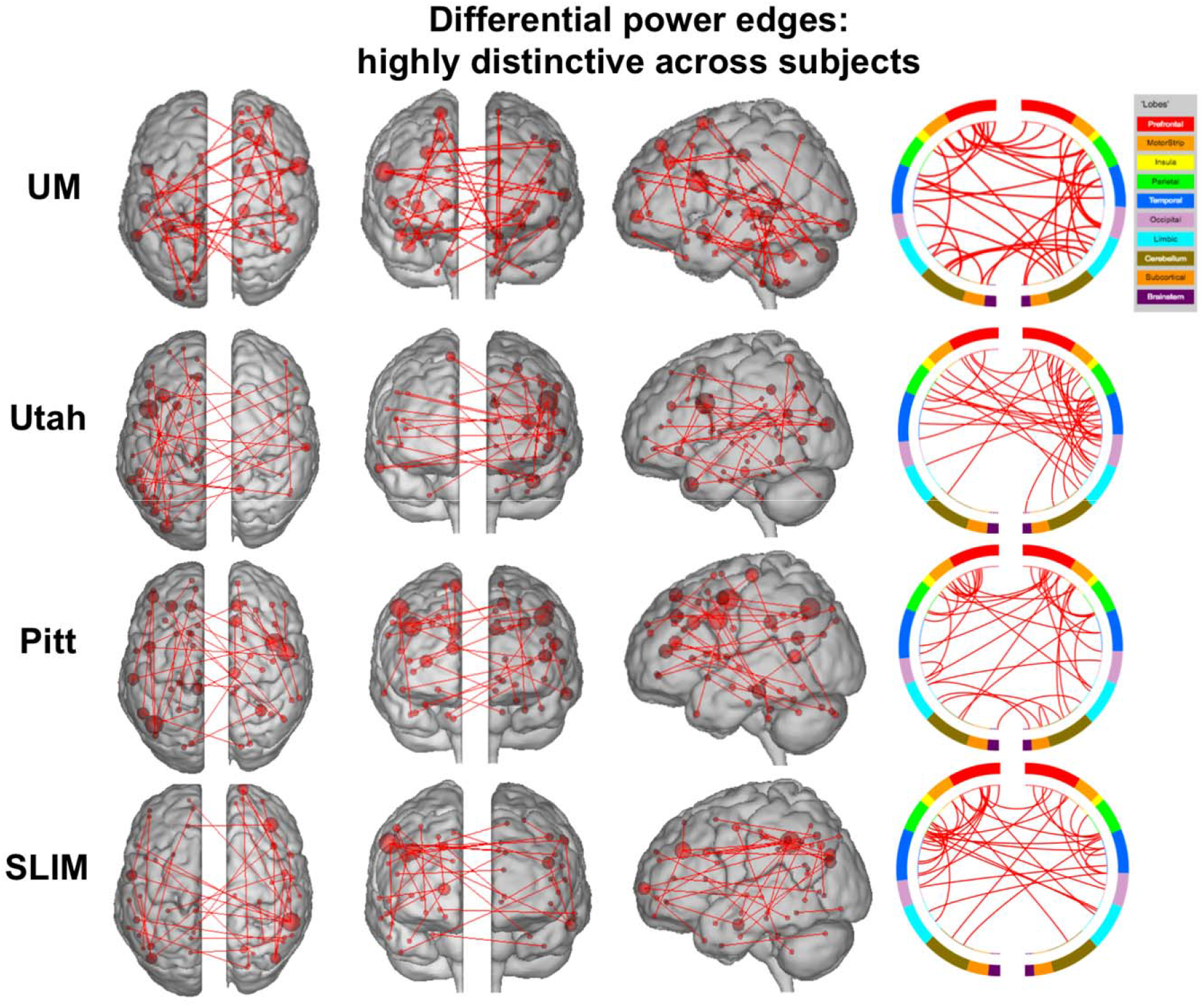
Results of edge-based analyses – differential power (highly discriminative) edges in low motion subjects. The edges shown here were in the top 99.9 percentile of highly unique edges among subjects – these edges were most helpful in an identification. Each row corresponds to results from a different dataset. For a given row, in the three leftmost images of the brain, the red lines indicate edges connecting the red spheres, representing nodes. Nodes are sized according to degree, the number of edges connected to that node. For a given row, on the right, the same nodes and edges are visualized on a circle plot, in which nodes are grouped according to anatomic location. The top of the circle represents anterior; the bottom, posterior. The left half of the circle plot corresponds to the left hemisphere of the brain. A legend indicating the approximate anatomic ‘lobe’ is shown to the top right of the figure. Results are shown for session 1 – session 2. Note that we considered Utah session 2a as session 2; for UM we considered session1a as session 1 and session 2a as session 2.

**Figure 8.**
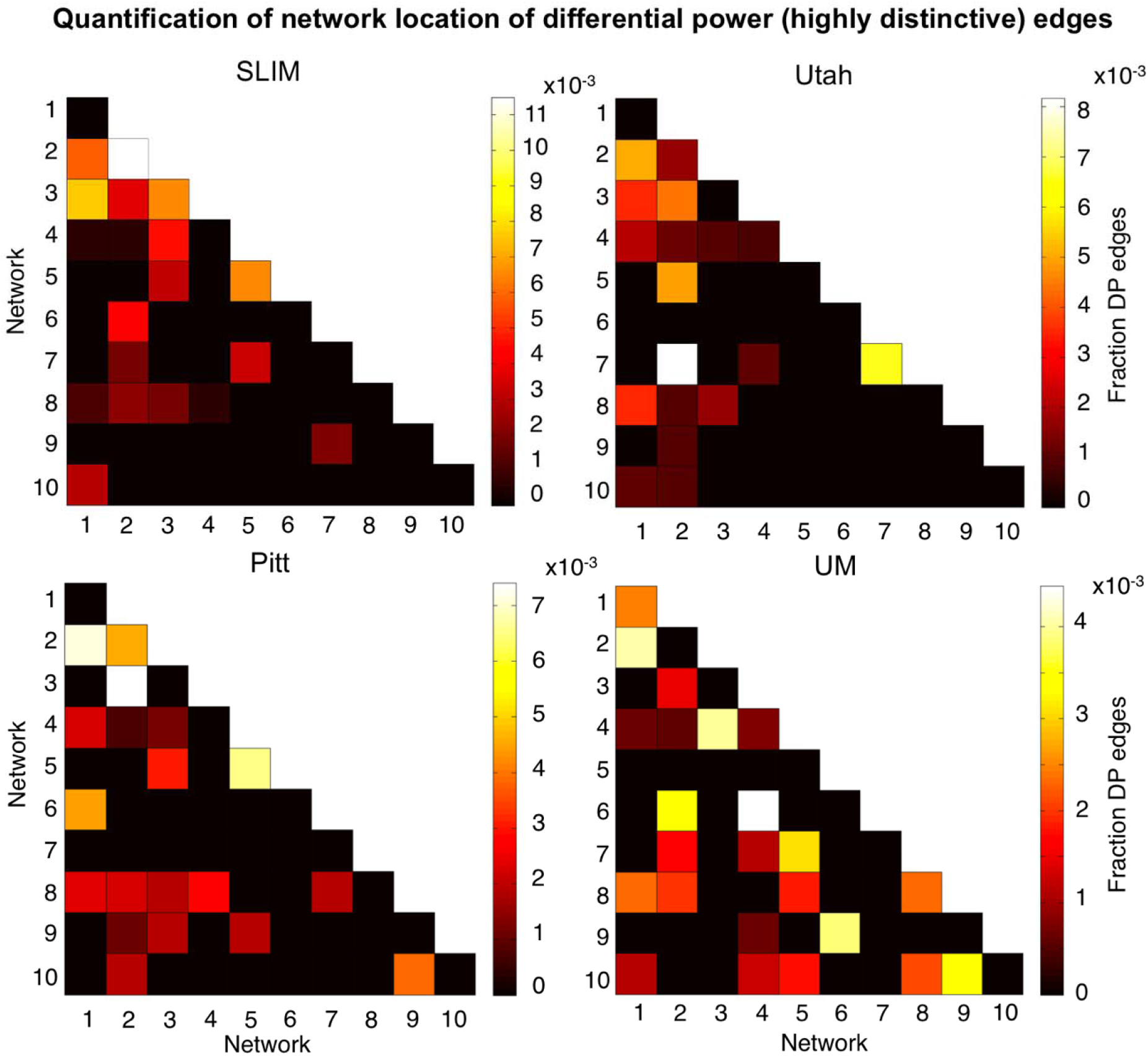
Network representation of differential power (highly unique) edges in low motion subjects. Here we are quantifying the network location of the data shown in Figure 7 (i.e. these are the 99.9 percentile most unique edges). Hotter colors indicate more edges are in the network pair; if an element in the matrix is black it indicates no edges were in this network pair. Results are normalized by network size. Data are shown for session 1 – session 2. Note that the color bars are not necessarily the same among all datasets. Note also that we considered Utah session 2a as session 2; for UM we considered session1a as session 1 and session 2a as session 2.

To quantify the extent to which individual edges do not contribute to subject uniqueness, we calculated the group consistency measure, which quantifies edges that are highly consistent within a single subject and across all subjects in a dataset. Because they are highly consistent for all subjects in a dataset, such edges do not discriminate between individuals. For visualization we show the anatomical locations of group consistency edges in the 99.9 percentile; these edges tended to link cross-hemispheric homologs (Figure 9). This pattern of connecting to cross-hemispheric homologs was stable across a range of thresholds (Supplemental Figure 14). Quantifying network representation revealed that the majority of these edges tended to be located in visual networks (networks 5 and 6), as well as the cerebellum in UM (network 10; Figure 10). Group consistency results were stable when this analysis was repeated on all subjects in dataset (Supplemental Figure 15 and 16).

**Figure 9.**
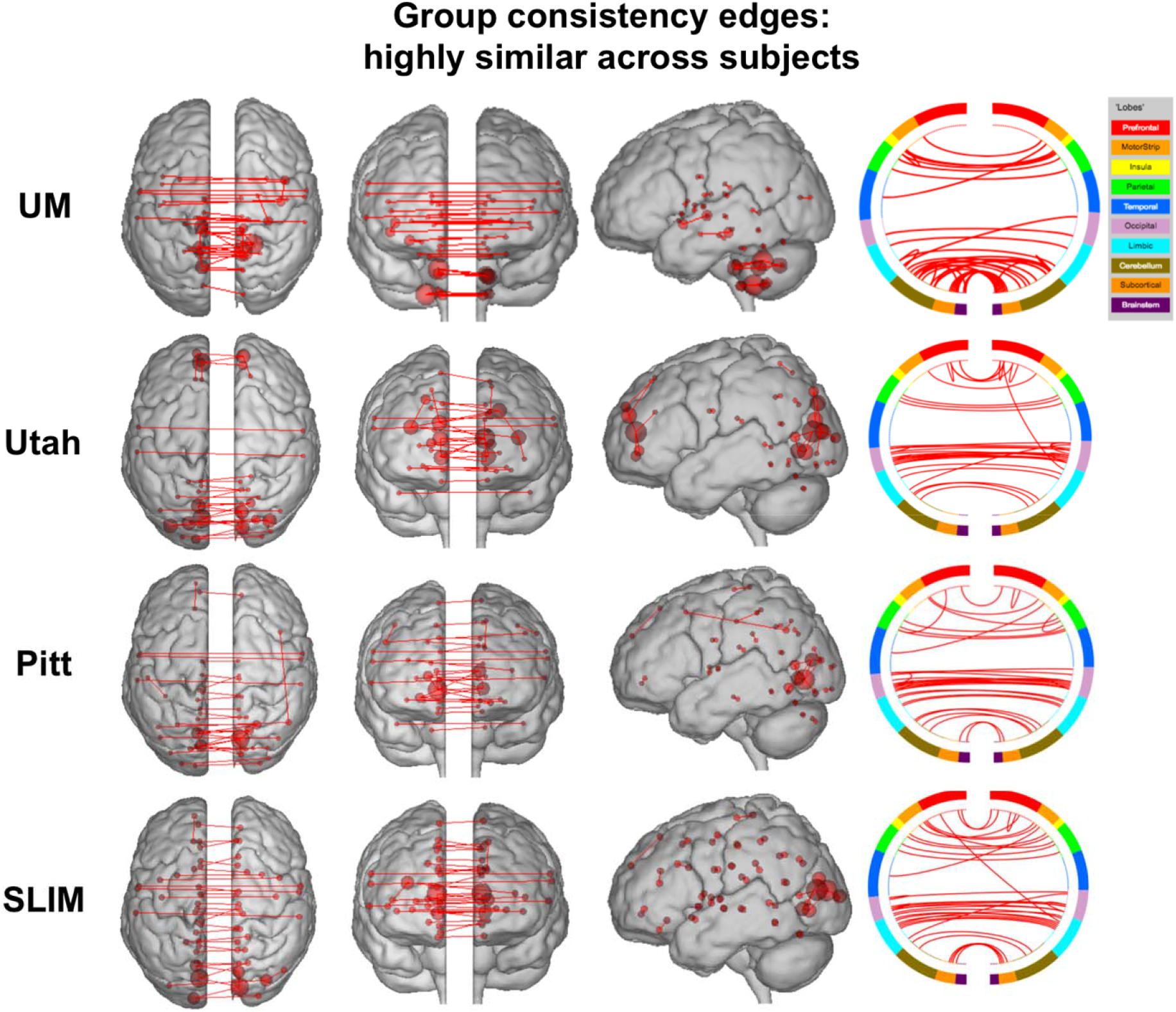
Results of edge-based analyses – group consistency (highly similar) edges in low motion subjects. The edges shown here were in the top 99.9 percentile of highly similar edges among subjects – these edges were least helpful in an identification. Each row corresponds to results from a different dataset. For a given row, in the three leftmost images of the brain, the red lines indicate edges connecting the red spheres, representing nodes. Nodes are sized according to degree, the number of edges connected to that node. For a given row, on the right, the same nodes and edges are visualized on a circle plot, in which nodes are grouped according to anatomic location. The top of the circle represents anterior; the bottom, posterior. The left half of the circle plot corresponds to the left hemisphere of the brain. A legend indicating the approximate anatomic ‘lobe’ is shown to the top right of the figure. Results are shown for session 1 – session 2. Note that we considered Utah session 2a as session 2; for UM we considered session1a as session 1 and session 2a as session 2.

**Figure 10.**
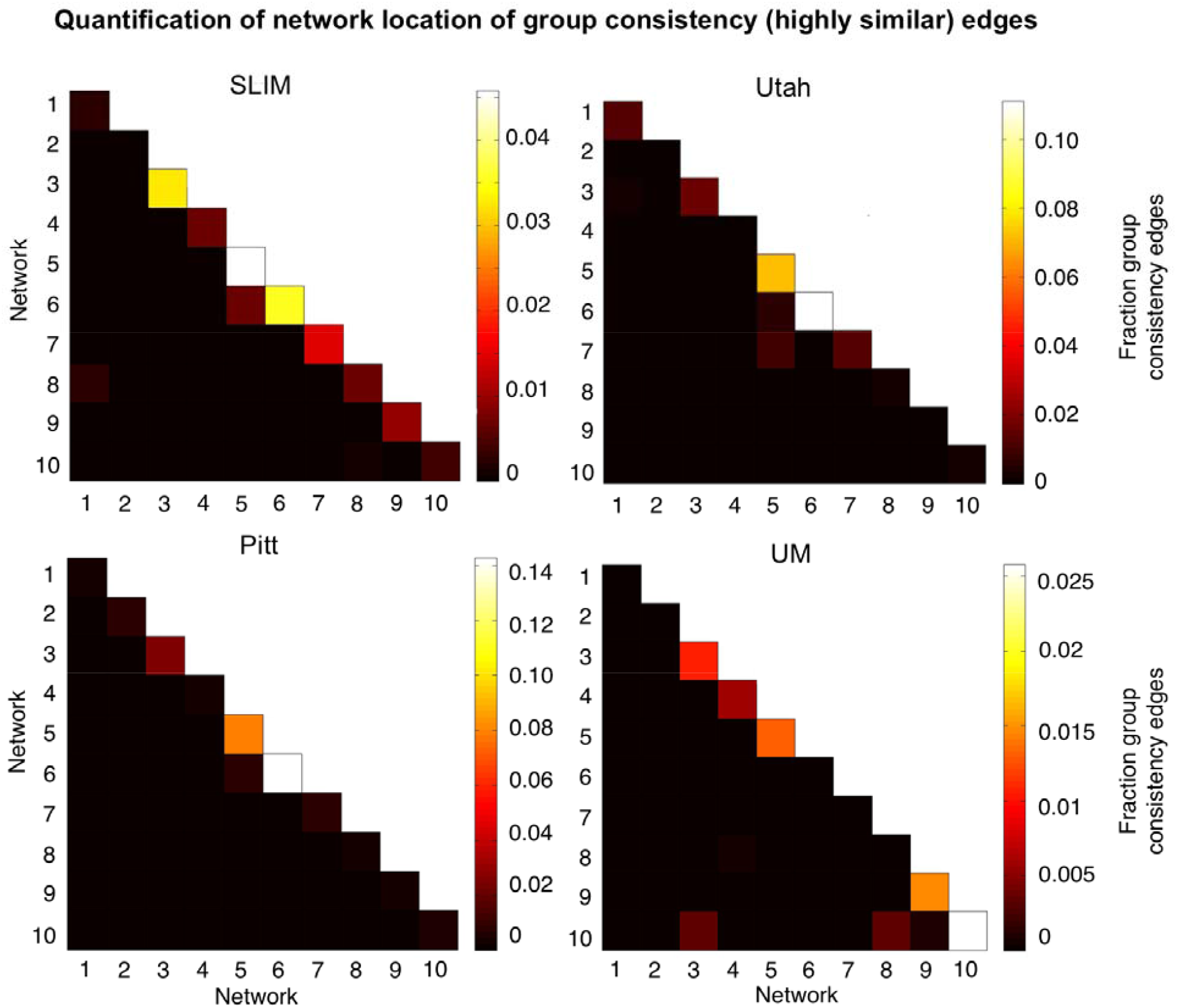
Network representation of group consistency (highly similar) edges in low motion subjects. Here we are quantifying the network location of the data shown in Figure 9 (i.e. these are the 99.9 percentile most similar edges). Hotter colors indicate more edges are in the network pair; if an element in the matrix is black it indicates no edges were in this network pair. Results are shown for session 1 – session 2. Note that the color bars are not necessarily the same for all datasets. Note also that we considered Utah session 2a as session 2; for UM we considered session1a as session 1 and session 2a as session 2.

## Discussion

In this work, we set out to assess the stability of the functional connectome over longer periods of time, as well as the extent to which unique networks over the course of days (i.e. medial frontal and frontal parietal networks; Finn et al., 2015) remain unique across months to years. Utilizing connectome-based identification in four longitudinal resting-state fMRI datasets, we determined that individual differences in the functional connectome are indeed stable across at least three years. Further, the networks driving individual uniqueness across days also drive uniqueness across months to years. By utilizing samples comprised of younger participants (children and adolescents) as well as older adults, our results also demonstrate that even in periods of significant brain changes, individuals retain a unique signature of functional connectivity patterns. Hence, consistent with other recent work (Finn et al., 2015; Gordon et al., 2017b; Gratton et al., 2018; Laumann et al., 2015; Poldrack et al., 2015), we find that individual differences are a stable feature of brain organization revealed by functional connectivity analyses.

### Replication and open science

An important aspect of the scientific process is reliability and reproducibility, and practical steps have been proposed in the context of fMRI (Nichols et al., 2017; Poldrack et al., 2017). Especially in a field like neuroimaging, which has largely failed to make an impact on clinical practice, studies are needed that assess the replicability and generalizability of findings (Nichols et al., 2017; Poldrack et al., 2017). Given the advent of the open data movement in neuroimaging (Poldrack et al., 2017; Poldrack et al., 2013; Poldrack and Gorgolewski, 2014; Zuo et al., 2014), there are multiple open source datasets readily available to assess the reproducibility of findings. Here, we leveraged these resources to use four different datasets, with different sample characteristics, to address the stability of functional connectivity patterns. Given the need to utilize test-retest datasets, we would not have been able to conduct the analyses reported here if it were not for open source data. Using these resources allowed us to replicate the findings of previous reports (Amico and Goni, 2018; Biazoli et al., 2017; Finn et al., 2017; Finn et al., 2015; Horien et al., 2018; Kaufmann et al., 2018; Kaufmann et al., 2017; Miranda-Dominguez et al., 2014; Miranda-Dominguez et al., 2018; Noble et al., 2017; Vanderwal et al., 2017; Waller et al., 2017) and add to the growing body of literature in connectome-based identification studies by extending this work, specifically focusing on identification across longer time frames. Hence, our overall approach speaks to the power of open data in neuroimaging, both in confirming and advancing initial results.

### How can a network be “stable” but undergo developmental changes?

It has previously been shown that networks comprised of nodes in frontal and parietal association cortices are important for defining individual uniqueness across a shorter time frame (Finn et al., 2017; Finn et al., 2015; Kaufmann et al., 2017; Miranda-Dominguez et al., 2014; Vanderwal et al., 2017; Waller et al., 2017). Our new findings extend these results by showing that these same areas are stable over longer time frames. While our findings are consistent with other work highlighting the degree to which these regions exhibit differences between subjects (Finn et al., 2015; Gordon et al., 2017a; Gratton et al., 2018; Miranda-Dominguez et al., 2014; Mueller et al., 2013), our results are intriguing given that these same regions undergo changes in children and adolescents (Giedd et al., 1999; Gogtay et al., 2004; Gu et al., 2015; Kaufmann et al., 2017; Vasa et al., 2018) as well as in older adults (Chan et al., 2014; Damoiseaux, 2017; Ng et al., 2016). One possible explanation lies in the individualized methods used here. The connectome-based ID approach does not require a participant have precisely the same connectivity matrix from scan-to-scan; all that is needed for a correct ID is that a subject must look more like her/himself than anyone else. Hence, the high ID rates obtained here add a layer of nuance to the development literature: children and adolescents (and older adults) undergo neurodevelopmental changes leading to differences in functional connectivity measures due to age, but these changes occur in an individual-specific way. That is, despite brain changes, individuals still tend to look like themselves across development.

### A word about motion

Given the effect of in-scanner movement on estimates of functional connectivity (Power et al., 2015; Satterthwaite et al., 2012), motion is an important variable to consider. Motion has also been demonstrated to exhibit high test-retest reliability (Zuo and Xing, 2014), potentially complicating identification. We therefore undertook numerous steps to address movement in our preprocessing and analysis pipelines, and we observed that identification rates were high independent of motion. We also tested identification rates using motion distribution vectors, which capture subject-specific movement, and found it was not possible to identify subjects. This result is consistent with earlier studies (Amico and Goni, 2018; Finn et al., 2017; Finn et al., 2015; Horien et al., 2018; Vanderwal et al., 2017), and reinforces that the results obtained here are due to differences in the functional connectivity data as opposed to a movement-related artifact.

### Individual differences and individual parcellations

We applied a group parcellation to our resting-state data, as have all connectome-based identification studies to this point. Recent work has indicated that individuals exhibit meaningful differences in the spatial topography of networks (e.g. differences in spatial topography are predictive of phenotypic measures; Kong et al., 2018), and that these differences can confound estimates of connectivity when a group parcellation is used (Bijsterbosch et al., 2018). Thus, it is possible that individual differences in spatial topography are driving subject identification (along with, or in place of, differences in functional connectivity). A potential way to circumvent this confound is using individualized parcellations (e.g. Braga and Buckner, 2017; Gordon et al., 2017b; Kong et al., 2018; Laumann et al., 2015; Salehi et al., 2017; Wang et al., 2015). Given that our goal was confirming and extending work using group parcellations, we did not individualize parcellations here. Further, given that substantial amounts of data have sometimes been required to establish accurate individual differences (Laumann et al., 2015; Gordon et al., 2017; Braga and Buckner, 2017), how best to use and validate these methods with “typical” scan times (i.e. the resting-state scans used here with 5-8 minutes per scan) is unclear, especially considering that little individualized parcellation work has been conducted in younger children and older adults, a majority of the participants in this study. Applying individualized parcellation techniques in these populations could be a fruitful area of future research, as could examining the stability of network topography more generally.

### Relevance to phenotypic measures

One of the key goals of precision neuroscience is relating individual differences in brain data to differences in phenotypic/clinical data. Indeed, this was an important component of the original identification study (Finn et al., 2015) and much exciting work has since been conducted in this area using a variety of methods (Beaty et al., 2018; Drysdale et al., 2017; Greene et al., 2018; Hearne et al., 2016; Hsu et al., 2018; Jangraw et al., 2018; Kaufmann et al., 2017; Kong et al., 2018; Rosenberg et al., 2016; Shen et al., 2017). We were unable to add to this growing literature, because only the SLIM dataset included behavioral measures, but future work could determine if the stability of single-subject fMRI data allows predictions to be made regarding behavior at longer time frames, similar to how previous investigators have used functional connectivity measures to predict a behavioral or psychiatric score in cross-sectional studies.

### Limitations and future directions

We only examined resting-state data here; hence, the extent to which task-based connectivity patterns remain stable is unclear. Other studies have shown that identification rates are highest across days when subjects are completing a task in the scanner (possibly by increasing subject-specific signal-to-noise and augmenting unique patterns of functional connectivity; (Finn et al., 2017; Vanderwal et al., 2017), so identifiability could be investigated across longer time frames when subjects are completing task-based scans to see if a similar pattern holds. Task-based connectivity studies have recently been shown to lead to better modeling of the link between brain and behavior, again emphasizing how individual traits can be amplified by performance of a task (Greene et al., 2018). Another limitation is that we were unable to compare ID rates across samples given the numerous differences among the datasets (eyes open versus eyes closed resting-state runs, differences in acquisition sequences, differences in the length of times between scans, etc.). Thus, questions about how identifiability varies as a function of age in longitudinal samples await further investigation.

A number of other questions (Finn et al., 2015; Finn and Constable, 2016) remain open for study. For example, is an individual’s functional connectome unique from birth (or even in utero), or is the distinctiveness something that develops at a later age? If so, what is the developmental trajectory? The stability of the connectome across even longer time frames is unknown (i.e. across decades) and could be investigated. Finally, it is also unclear if an individual’s connectome ever becomes unidentifiable from scan-to-scan. Does identifiability degrade at a certain age? If so, how does this relate to measures of behavior and cognition?

### Conclusions

In sum, we have shown that subjects have unique and stable functional connectomes over the span of years, and that medial frontal and frontoparietal networks remain important in defining individual uniqueness over these long time scales. Leveraging the stability of functional connectome data to generate meaningful models related to phenotypic or clinical variables remains an important goal for this field.

## Acknowledgements

This work was supported by a Medical Scientist Training Program training grant (NIH/NIGMS T32GM007205; C.H.)

